# GenPipes: an open-source framework for distributed and scalable genomic analyses

**DOI:** 10.1101/459552

**Authors:** Mathieu Bourgey, Rola Dali, Robert Eveleigh, Kuang Chung Chen, Louis Letourneau, Joel Fillon, Marc Michaud, Maxime Caron, Johanna Sandoval, Francois Lefebvre, Gary Leveque, Eloi Mercier, David Bujold, Pascale Marquis, Patrick Tran Van, David Morais, Julien Tremblay, Xiaojian Shao, Edouard Henrion, Emmanuel Gonzalez, Pierre-Olivier Quirion, Bryan Caron, Guillaume Bourque

## Abstract

With the decreasing cost of sequencing and the rapid developments in genomics technologies and protocols, the need for validated bioinformatics software that enables efficient large-scale data processing is growing. Here we present GenPipes, a flexible Python-based framework that facilitates the development and deployment of multi-step workflows optimized for High Performance Computing clusters and the cloud. GenPipes already implements 12 validated and scalable pipelines for various genomics applications, including RNA-Seq, ChIP-Seq, DNA-Seq, Methyl-Seq, Hi-C, capture Hi-C, metagenomics and PacBio long read assembly. The software is available under a GPLv3 open source license and is continuously updated to follow recent advances in genomics and bioinformatics. The framework has been already configured on several servers and a docker image is also available to facilitate additional installations. In summary, GenPipes offers genomic researchers a simple method to analyze different types of data, customizable to their needs and resources, as well as the flexibility to create their own workflows.

## INTRODUCTION

Sequencing has become an indispensable tool in our quest to understand biological processes. Moreover, facilitated by a significant decline in overall costs, new technologies and experimental protocols are being developed at a fast pace. This has resulted in massive amounts of sequencing data being produced and deposited in various public archives. For instance, a number of national initiatives, such as *Genomics England* and *All of US*, plan to sequence hundreds of thousands of individual genomes in an effort to further develop precision medicine. Similarly, a number of large initiatives, such as ENCODE [1] and the International Human Epigenome Consortium (IHEC) [2], plan to generate thousands of epigenomics datasets to better understand gene regulation in normal and disease processes. Despite this rapid progress in sequencing, genomics technologies and available datasets, processing and analyses have struggled to keep up. Indeed, the need for robust, open-source and scalable bioinformatics pipelines has become a major bottleneck for genomics [3].

Available bioinformatics tools for genomic data can be categorized into three different groups: 1) analysis platforms/workbenches, 2) workflow management systems (WMS)/frameworks, and 3) individual analysis pipelines/workflows. Platforms of the first type, like Galaxy [4] or DNA Nexus [5], provide a full workbench for data upload and storage, and are accompanied with a set of available tools. While they provide fast and easy user services, such tools can be inconvenient for large scale projects. In the second type, WMSs such as Snakemake [6], Nextflow [7], BPipe [8], BigDataScript [9] and declarative workflow description languages, such as CWL or WDL are dedicated to providing a customizable framework to build bioinformatics pipelines. Such solutions are flexible and can help in pipeline implementation but rarely provide robust pre-built pipelines which are ready for production analysis. Finally, tools of the third type are individual analysis pipelines for various applications that have been validated and published. These are useful for specific applications but can sometimes be challenging to implement, difficult to modify or scale-up. They have also rarely been tested on multiple computing infrastructures.

Here we present GenPipes, an open-source, Python-based WMS for pipeline development. As part of its implementation, GenPipes includes a set of high-quality, standardized analysis pipelines, designed for High Performance Computing (HPC) resources and cloud environments. GenPipes’ WMS and pipelines have been tested, benchmarked and used extensively over the past four years. GenPipes is continuously updated and is configured on several different HPC clusters with different properties. By combining both WMS and extensively validated End-to-End analysis workflows, GenPipes offers turnkey analyses for a wide range of bioinformatics applications in the genomics field while also enabling flexible and robust extensions.

## MATERIAL AND METHODS

### Overview of the GenPipes Framework

GenPipes is an object-oriented framework consisting of Python scripts and libraries which create a list of jobs to be launched as Bash commands (Figure 1). There are four main objects that manage the different components of the analysis workflow, namely, *Pipeline*, *Step*, *Job and Scheduler*. The main object is the “*Pipeline*” object which controls the workflow of the analysis. Each specific analysis workflow is thus defined as a specific *Pipeline* object. *Pipeline* objects can inherit from one another. The *Pipeline* object defines the flow of the analysis by calling specific *“Step”* objects. The *Pipeline* instance could call all steps implemented in a pipeline or only a set of steps selected by the user. Each step of a pipeline is a unit block that encapsulates a part of the analysis (e.g., trimming or alignment). The *Step* object is a central unit object which corresponds to a specific analysis task. The execution of the task is directly managed by the code defined in each *Step* instance; some steps may execute their task on each sample individually while other steps execute their task using all the samples collectively. The main purpose of the *Step* object is to generate a list of *“Job”* objects which correspond to the consecutive execution of single tasks. The *Job* object defines the commands that will be submitted to the system. It contains all the elements needed to execute the commands, such as input files, modules to be loaded, as well as job dependencies and temporary files. Each *Job* object will be submitted to the system using a specific “*Scheduler*” object. The *Scheduler* object creates execution commands that are compatible with the user’s computing system. Four different *Scheduler* objects have already been implemented (PBS, SLURM, Batch and Daemon), see below.

**Figure 1.**
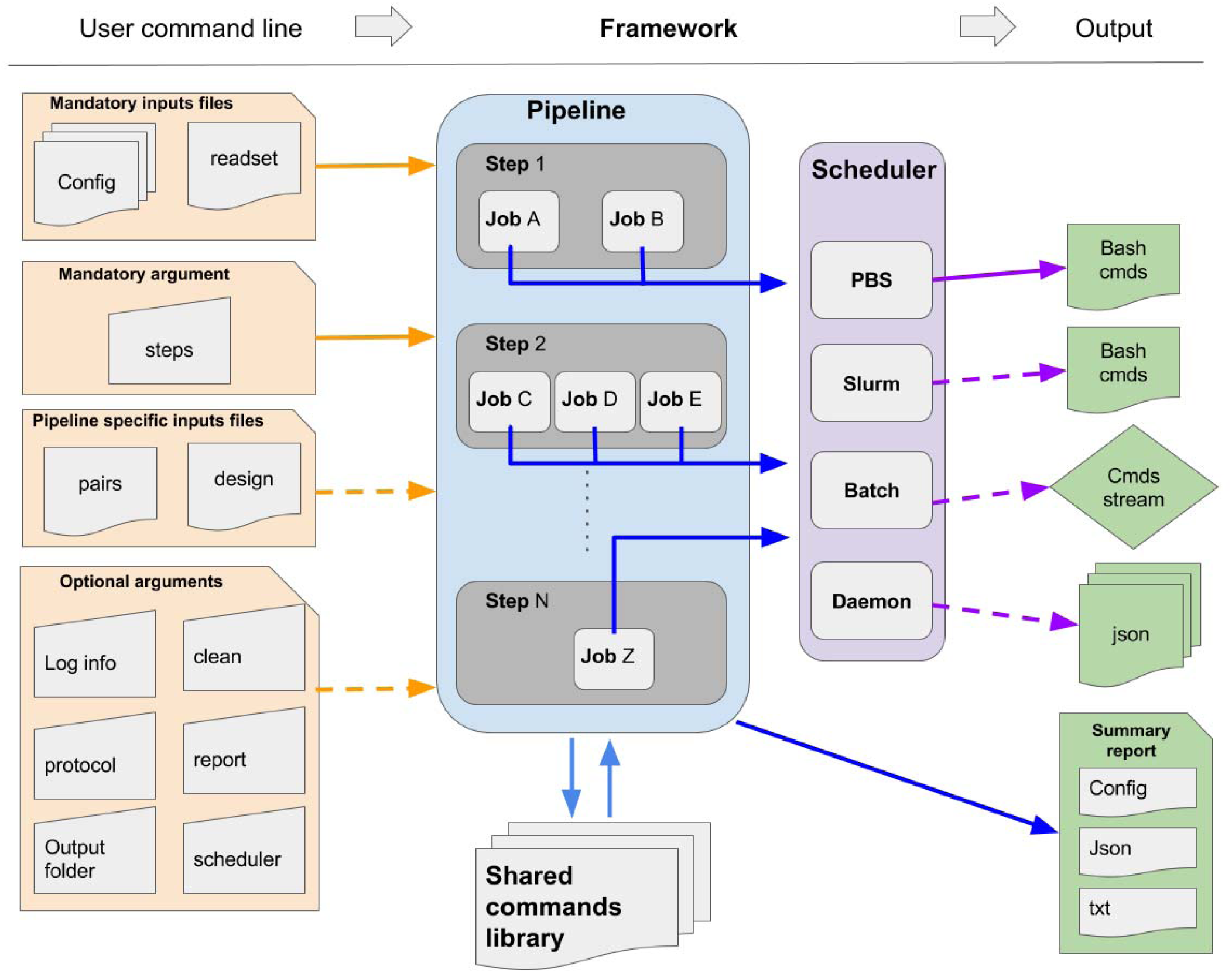
General workflow of GenPipes. Diagram showing how the information flows from the user command line input through the 4 different objects *(Pipeline, Step, Job and Scheduler)* in order to generate system specific executable outputs.

GenPipes’ object-oriented framework simplifies the development of new features and its adaptation to new systems; new workflows can be created by implementing a *Pipeline* object which inherits features and steps from other existing *Pipeline* objects. Similarly, deploying GenPipes on a new system may only require the development of the corresponding *Scheduler* object along with specific configuration files. GenPipes’ command execution details have been implemented using a shared library system which allows the modification of tasks by simply adjusting input parameters. This simplifies code maintenance and makes changes in software versions consistent across all pipelines.

### Freely distributed and pre-installed on a number of HPC resources

GenPipes is an open-source framework freely distributed and open for external contributions from the developer community. GenPipes can be installed from scratch on any Linux cluster supporting Python 2.7 by following the available instructions (https://bitbucket.org/mugqic/genpipes/src/master/). GenPipes can also be used via a Docker image which simplifies the setup process and can be used on a range of platforms, including cloud platforms. This allows system-wide installations, as well as local user installations via the Docker image without needing special permissions.

Through a partnership with the Compute Canada consortium (https://www.computecanada.ca), the pipelines and third-party tools have also been configured on 6 different Compute Canada HPC centers. It allows any Canadian researcher to use GenPipes along with the needed computing resources by simply applying to the consortium [10]. To ensure consistency of pipeline versions and used dependencies (such as genome references and annotation files) and to avoid discrepancy between compute sites, pipeline setup has been centralized to one location which is then distributed on a real-time shared file system: the CERN Virtual Machine File System [11].

### Running GenPipes

GenPipes is a command line tool. Its use has been simplified to accommodate general users. A full tutorial is available [12]. Briefly, to launch GenPipes, the following is needed:

- A readset file that contains information about the samples, indicated using the flag “-r”.
- Configuration/ini files that contain parameters related to the cluster and the third-party tools, indicated using the flag “-c”.
- The specific steps to be executed, indicated by the flag “-s”.

The generic command to run GenPipes is: <pipeline>.py -c myConfigurationFile -r myReadSetFile -s 1-X > Commands.txt && bash Commands.txt

Where <pipeline> can be any of the 12 available pipelines and X is the step number desired. Commands.txt contains the commands that the system will execute.

Pipelines that conduct sample comparisons, like ChIP-Seq and RNA-Seq, require a design file that describes each contrast. Design files are indicated by the flag “-d”. The tumour_pair pipeline requires normal-tumour pairing information provided in a standard CSV file using the “-p” option. For more information on the design file and the content of each file type, please consult the GenPipes tutorial and the online documentation.

When the GenPipes command is launched, required modules and files will be searched for and validated. If all required modules and files are found, the analysis commands will be produced. GenPipes will create a directed acyclic graph (DAG) that defines job dependency based on input and output of each step. For a representation of the DAG of each pipeline, refer to supplementary figures S1-14. Once launched, the jobs are sent to the scheduler and queued. As jobs complete successfully, their dependent jobs are released by the scheduler to run. If a job fails, all its dependent jobs are terminated and an email notification is sent to the user. When GenPipes is re-run, it will detect which steps have successfully completed, as described in section ‘Smart relaunch features’, and skip them but will create the command script for the jobs that were not completed successfully. To force the entire command generation, despite successful completion, the “-f” option should be added.

## RESULTS

GenPipes was first released in 2014. Since then, it has grown to implement 12 pipelines and is currently installed and maintained on 13 different clusters (Figure 2a-b). GenPipes has been actively used for the last four years to quality control and analyze thousands of samples each year (Figure 2c). It has also been used to analyze data for several large-scale projects such as IHEC [2] and eFORGE [13].

**Figure 2.**
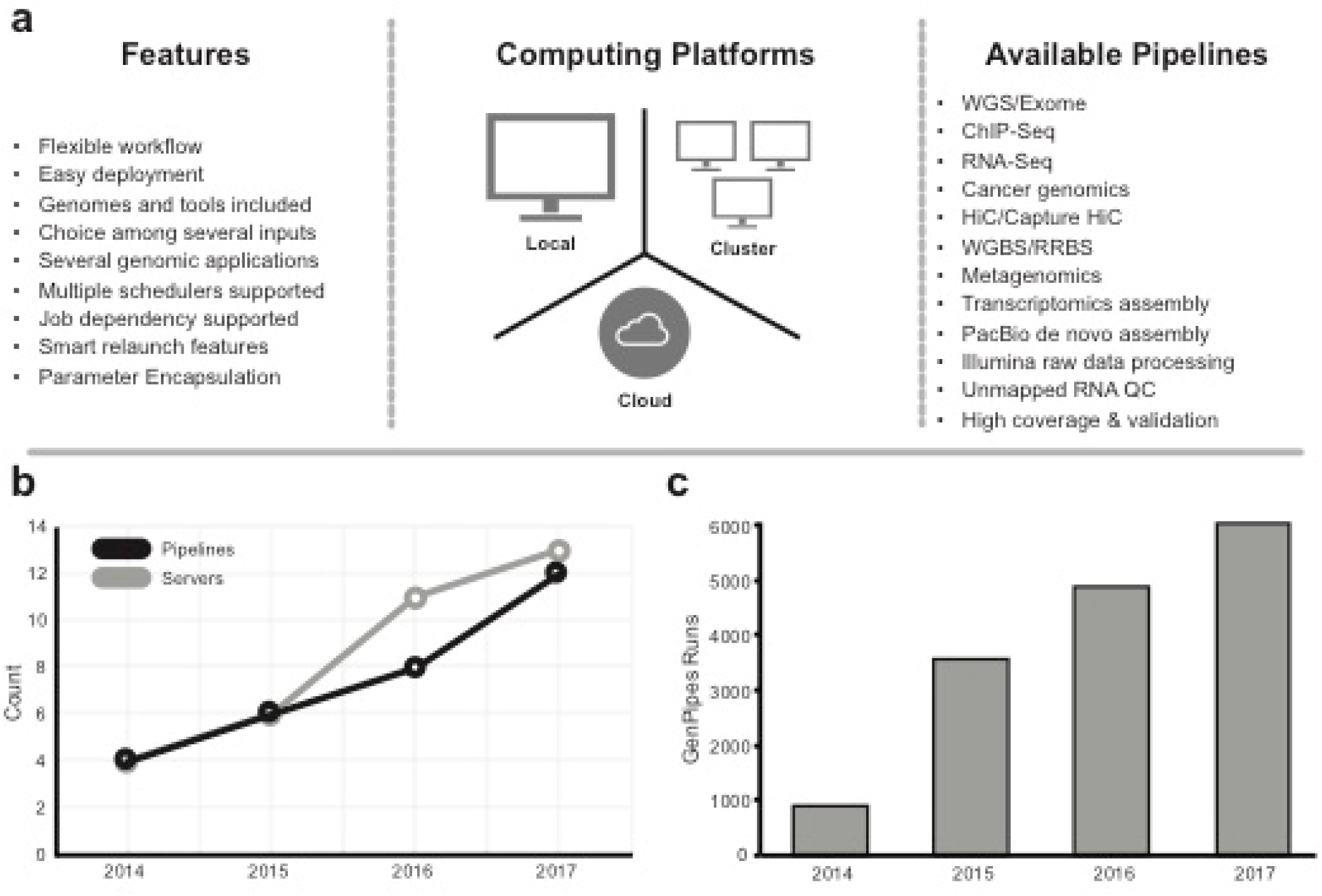
GenPipes properties. GenPipes’ properties and growth. (a) Diagram showing GenPipes’ features, compatible computing platforms and available pipelines. (b) GenPipes’ available pipelines and maintained servers since the release of GenPipes in 2014. (c) Bar plot showing the number of GenPipes runs per year since its release.

### Key features of GenPipes

GenPipes’ framework has been optimized to facilitate large scale data analysis. Several features make this possible (Figure 2a):

### Multiple schedulers

GenPipes is optimized for HPC processing. It can currently accommodate four different types of schedulers:

- *PBSScheduler* creates a batch script that is compatible with a PBS (TORQUE) system.
- *SLURMscheduler* creates a batch script that is compatible with a SLURM system.
- *BatchScheduler* creates a batch script which contains all the instructions to run all the jobs one after the other.
- *DaemonScheduler* creates a log of the pipeline command in a JSON file.

### Job dependencies

In order to minimize the overall analysis time, GenPipes uses a dependency model based on input files, which is managed at the *Job* object level. A job does not need to wait for the completion of a previous step unless it is dependent on its output. Jobs thus become active and can be executed as soon as all their dependencies are met, regardless of the status of previous jobs or of other samples. Thus, when a pipeline is run on multiple samples, it creates several dependency paths, one per sample, each of which completes at its own pace.

### Smart relaunch features

Large scale data analysis is subject to failure which could occur due to system failure (e.g. power outage, system reboot, etc…) or user failure (errors in set parameters, or resources). To limit the micromanagement and time required to relaunch the pipeline from scratch, GenPipes includes a system of reporting which provides the status of every job in the analysis in order to facilitate the detection of jobs which have failed. Additionally, a relaunch system is implemented which allows restarting the analysis at the exact state before the failure. The relaunch system uses two features: md5sum hash and time stamps. When GenPipes is launched, a md5sum hash is produced for each command. Upon relaunch following a failure, the newly produced hash is compared to that of the completed job to detect changes in the commands. If the hashes are different, the job is relaunched. To detect updates in input files, GenPipes compares the time stamp on the input and output files of already completed jobs. If the date stamp on the input files is more recent than that on the output files then the job is relaunched. If neither the hash code nor the time stamp flag the job to be relaunched then it is considered complete and up-to-date and it will be skipped in the pipeline restart process.

### Configuration files

Running large-scale analyses requires a very large number of parameters to be set. GenPipes implements a superposed configuration system to reduce the time required to set-up or modify parameters needed during the analysis. Configuration files, also referred to as “ini” files, are provided among the arguments of the GenPipes command. These files follow the standard INI format, which was selected for its readability and ease of use by non-expert users. Each pipeline reads all configuration files, one after the other, based on a user defined order. The order is of major importance as the system will overwrite a parameter each time it is specified in a new ini file. The system allows the use of the default configuration files provided in GenPipes alone or in combination with user specific configuration files. Configuration files provided with GenPipes are the result of years of experience along with intensive benchmarking. Additionally, several configuration files adjusted for different compute systems or different model organisms are available. The main advantage of this system is to reduce the users’ task; only parameters that need to be modified (e.g system parameters, genomic resources, user specific parameters) have to be adjusted during the set-up phase of the analysis. To track and enable reproducibility, GenPipes always outputs a file containing the final list of parameters used for the analysis.

### Choice among multiple inputs

GenPipes represents a series of *Step* objects that are interdependent based on inputs and outputs. Many of the pipeline steps implemented in GenPipes, represent filtering, manipulation or modification of specific genomics files share common formats (e.g. bam, fastq, vcf). To ensure more flexibility in the analysis, a system of ordered list to be interpreted as input files is used. For a given *Step*, each *Job* can be given a series of inputs. The *Job* will browse its list of possible inputs and will consider them based on the order in the list. The first input file found either on disk or in the overall output list will be chosen as input. The chosen input will determine the dependency of the *Job* to the other *Jobs* in the pipeline. This system is really flexible and allows users to skip specific steps in the pipeline if they consider them unnecessary.

### Customizable workflows

Despite the benchmarking and testing made on the standard analysis procedures implemented in GenPipes, some users may be interested in modifying pipelines. In order to make GenPipes more flexible, a *protocol* system is used. The system allows the implementation of different workflows into a single *Pipeline* object. As a result, one can replace specific steps by other user specific ones. In that case, the user will only need to implement these new Steps and define an additional protocol which will use part of the initial Steps and the newly developed ones. As an example, this has been used to incorporate the Hi-C analysis workflow and the capture Hi-C analysis workflow into GenPipes’ hicseq pipeline. A flag (-t hic or -t capture) can be used to specify the workflow to be executed. This system has been developed to reduce the amount of work for external users that decide to contribute to code development and to limit the number of Pipeline objects to maintain. This will also allow us to provide multiple workflows per pipeline to appeal to different tool preferences in each field.

### Facilitating dependency installation

Genomic analyses require third party tools, as well as genome sequence files, annotation files and indices. GenPipes comes configured with a large set of reference genomes and their respective annotation files, as well as indices for most aligners. It also includes a large set of third party tools. If GenPipes is being installed from scratch on new clusters, automatic bash scripts that download all tools and genomes are included to ease the setup process. These scripts support local installations without the need for super-user privileges. Tools and dependencies are versioned and are loaded by GenPipes in a version-specific manner. This allows different pipelines to use different software versions based on need. It also allows retention of the same parameters and tools for any given project for reproducibility. GenPipes is also provided as a container version for which no dependency installation is required.

### Available workflows

GenPipes implements 12 standardized genomics workflows including: DNA-Seq, Tumour Analysis, RNA-Seq, de novo RNA-Seq, ChIP-Seq, PacBio assembly, Methyl-Seq, Hi-C, capture Hi-C, and Metagenomics (Figure 2c). All pipelines have been implemented following a robust design and development routine by following established gold standards standard operating protocols (SOP). Below we summarize GenPipes’ workflows; more details are available in the GenPipes documentation. For more details concerning computational resources used by each pipeline, refer to supplementary Table S1. All workflows accept a bam or a fastq file as input.

#### DNA-Seq Pipeline

DNA-Seq has been implemented optimizing the GATK best practices SOPs [14]. This procedure entails trimming raw reads derived from whole genome or exome data followed by alignment to a known reference, post alignment refinements and variant calling. Trimmed reads are aligned to a reference by the Burrows-Wheeler Aligner, bwa-mem [15]. Refinements of mismatches near indels and base qualities are performed using GATK indels realignment and base recalibration [14] to improve read quality post alignment. Processed reads are marked as fragment duplicates using picard mark duplicates [14] and SNP and small indels are identified using either GATK haplotype callers or samtools mpileup [16]. The Genome in a Bottle [17] dataset was used to select steps and parameters minimizing the false positive rate and maximizing the true positive variants to achieve a sensitivity of 99.7%, precision of 99.1% and F1-score of 99.4% (For more details, refer to Supplementary Materials). Finally, additional annotations are incorporated using dbNSFP [18] and/or Gemini [19XX] and quality control metrics are collected at various stages and visualized using MulitQC [20]. This pipeline has two different protocols, the default protocol based on the GATK variant caller, haplotype_caller, (“-t mugqic”, Figure 3) and one based on the mpileup/bcftools caller (“-t mpileup”, Figure S1). Another pipeline that is optimized for deep coverage samples, dnaseq_high_coverage, can be found in Figure S2.

**Figure 3.**
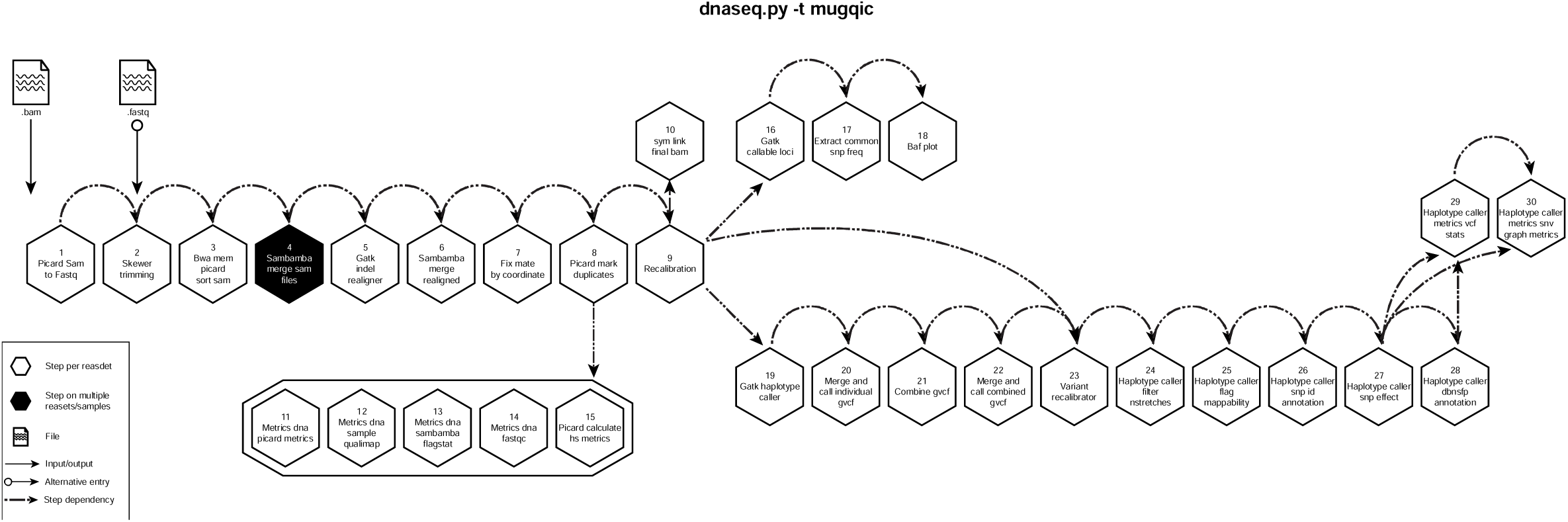
GenPipes DNASeq pipeline diagram. Schematic representation of GenPipes’ dnaseq.py pipeline. Hexagons represent steps in the pipeline. White hexagons represent steps that process input from a single sample, while black ones represent steps that process input from several samples. Arrows show step dependencies.

#### RNA-Seq Pipeline

This pipeline aligns reads with *STAR* [21] 2-passes mode, assembles transcripts with Cufflinks [22] and performs differential expression with *Cuffdiff* [23]. In parallel, gene-level expression is quantified using *htseq-count* [24], which produces raw read counts that are subsequently used for differential gene expression with both *DESeq* [25] and *edgeR* [26]. Several common quality metrics (rRNA content, expression saturation estimation etc.) are also calculated through the use of *RNA-SeQC* [27] and inhouse scripts. Gene Ontology terms are also tested for over-representation using *GOseq* [28]. Expressed short SNVs and indels calling is also performed by this pipeline, which optimizes GATK best practices to reach a sensitivity 92.8%, precision 87.7% and F1-score 90.1%. A schema of pipeline steps can be found in Figure S3. Another pipeline, rnaseq_light, based on Kallisto [29] and used for quick quality control can be found in Figure S4.

#### De-Novo RNASeq Pipeline

This pipeline is adapted from the Trinity-Trinotate suggested workflow [30] [31]. It reconstructs transcripts from short reads, predicts proteins and annotates leveraging several databases. Quantification is computed using RSEM and differential expression is tested in a manner identical to the RNA-seq pipeline. We observed that the default parameters of the Trinity suite are very conservative which could result in the loss of low-expressed but biologically relevant transcripts. In order to provide the most complete set of transcripts, the pipeline was designed with lower stringency during the assembly step in order to produce every possible transcript and not miss low expressed mRNA. A stringent filtration step is included afterward in order to provide a set of transcripts that make sense biologically. A schema of pipeline steps can be found in Figure S5.

#### ChIP-Seq Pipeline

The ChIP-Seq workflow is based on the ENCODE [1] workflow. It aligns reads using the BurrowsWheeler Aligner. It creates tag directories using Homer [32]. Peaks are called using MACS2 [33] and annotated using Homer. Binding motifs are also identified using Homer. Metrics are calculated based on IHEC requirements [34]. The ChIP-Seq pipeline can also be used for ATAC-Seq samples. However, we are developing a pipeline that is specific to ATAC-Seq. A schema of pipeline steps can be found in Figure S6.

#### The Tumour Analysis Pipeline

The Tumour Pair workflow inherits the bam processing protocol from DNA-seq implementation to retain the benchmarking optimizations but differs in alignment refinement and mutation identification by maximizing the information utilizing both tumour and normal samples together. The pipeline is based on an ensemble approach, which was optimized using both the DREAM3 challenge [35] and the CEPH mixture datasets to select the best combination of callers for both SNV and SV detection. For SNVs, multiple callers such as GATK mutect2, VarScan2 [36], bcftools and VarDict [37] were combined to achieve a sensitivity of 97.5%, precision of 98.8% and F1-score of 98.1% for variants found in 2 or more callers. Similarly, SVs were identified using multiple callers: DELLY [38], LUMPY [39], WHAM [40], CNVkit [41] and Svaba [42] and combined using MetaSV [43] to achieve a sensitivity of 84.6%, precision of 92.4% and F1-score of 88.3% for duplication variants found in the DREAM3 dataset (For more details, refer to Supplementary Material). The pipeline also integrates specific cancer tools to estimate tumour purity, tumour ploidy of sample pair normal-tumour. Additional annotations are incorporated to the SNV calls using dbNSFP [18] and/or Gemini [19] and quality control metrics were collected at various stages and visualized using MulitQC [20]. This pipeline has 3 protocols (sv, ensemble or fastpass). Schemas of pipeline steps for the three protocols can be found in Figures S7, 8 and 9.

#### Whole Genome Bisulfite Seq Pipeline (WGBS or Methyl-Seq)

The Methyl-Seq workflow is adapted from the Bismark pipeline [44]. It aligns paired-end reads with botiwe2 default mode. Duplicates are removed with Picard and methylation calls are extracted using bismark [44]. Wiggle tracks for both read coverage and methylation profile are generated for visualization. Variants calls can be extracted from the WGBS data directly using bisSNP [45]. Bisulfite conversion rates are estimated with lambda genome or from human non-CpG methylation directly. Several metrics based on IHEC requirements are also calculated. Methyl-Seq can also process capture data if provided with a capture bed file. A schema of pipeline steps can be found in Figure S10.

#### Hi-C Pipeline

The HiC-Seq workflow aligns reads using HiCUP [46]. It creates tag directories, produces interaction matrices, identifies compartments and significant interactions using Homer. It identifies Topologically Associating Domains using TopDom [47] and RobusTAD (bioRxiv 293175). It also creates “.hic” files using JuiceBox [48] and metrics reports using MultiQC [20]. The HiC-Seq workflow can also process capture Hi-C data with the flag “-t capture” using CHICAGO [49]. Schemas for the HiC and capture HiC protocols of this pipeline can be found in Figure S11 and Figure S12 respectively.

#### The Metagenomic Pipeline (rRNA gene amplification analysis)

This pipeline is based on the established Qiime procedure [50] for amplicon-based metagenomics. It assembles read pairs using FLASH [51], detects chimeras with uchime [52] and picks OTUs using vsearch [53]. OTUs are then aligned using PyNAST [54] and clustered with FastTree [55]. Standard diversity indices, taxonomical assignments and ordinations are then calculated and reported graphically. A schema of pipeline steps can be found in Figure S13.

#### The PacBio Pipeline

The PacBio whole genome assembly pipeline is built following the HGAP method [31], including additional features, such as base modification detection (https://github.com/PacificBiosciences/Bioinformatics-Training/wiki/Methylome-Analysis-Technical-Note) and genome circularization [56]. De novo assembly is performed using PacBio’s SMRT Link software (https://github.com/PacificBiosciences/SMRT-Link/wiki). Assembly contigs are generated using HGAP4. Alignments are then corrected and used as seeds by FALCON (https://github.com/PacificBiosciences/FALCON/wiki/) to create contigs. The resulting contigs are then polished and processed by “Arrow” (https://github.com/PacificBiosciences/GenomicConsensus) which ultimately generates high quality consensus sequences. An optional step allowing assembly circularization is integrated at the end of the pipeline. A schema of pipeline steps can be found in Figure S14.

### Comparison with other solutions for NGS analysis

Data collected for select tools, modified from Griffith & Griffith et al. [57] (Table 1), shows that GenPipes’ strength lies in its robust WMS that comes with one of the most diverse selection of analysis pipelines which have been thoroughly tested. The pipelines in the framework cover a wide range of sequencing applications (Figure 2a). The pipelines are end-to-end workflows running complete bioinformatics analyses. While many available pipelines conclude with a bam file or run limited post-bam analysis steps, the pipelines included in GenPipes are extensive, often having as many as 40 different steps that cover a wide range of post-bam processing. It is important to note that GenPipes, as well as several other WMSs, have community-supported pipelines, however, those have not been included in the comparison.

**Table 1.**
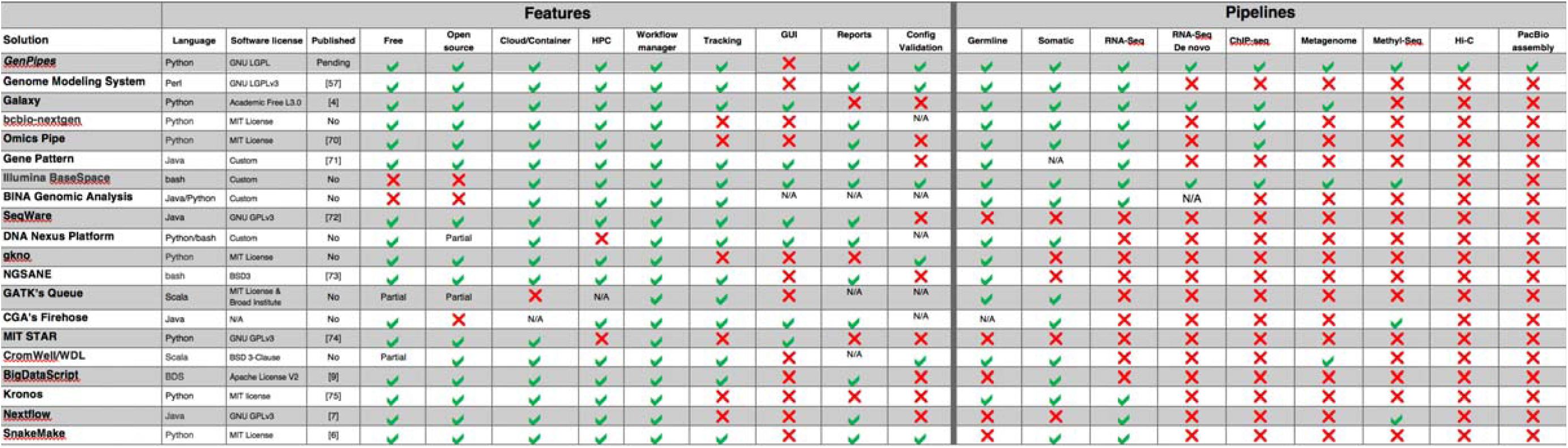
Comparison of available solutions for NGS analysis. Modified from Griffith & Griffith et al. [57].

GenPipes is compatible with HPC computing, as well as cloud computing [58] and includes a workflow manager that can be adapted to new systems. GenPipes also provides job status tracking through JSON files that can then be displayed on a web portal (an official portal for GenPipes will be released soon). GenPipes’ available pipelines facilitate bioinformatics processing, while the framework makes it flexible for modifications and new implementations.

GenPipes developers offer continuous support through a Google forum page [59] and a help desk email address. Since the release of version 2.0.0 in 2014, a community of users has run GenPipes to conduct approximately 3000 analyses processing around 100,000 samples (Figure 2b-c).

## DISCUSSION and CONCLUSION

GenPipes is a workflow management system that facilitates building robust genomic workflows. GenPipes is a unique solution which combines both a framework for development and end-to-end analysis pipelines for a very large set of genomics fields. The efficient framework for pipeline development has resulted in a broad community of developers with over 30 active branches and more than 10 forks of the GenPipes repository. GenPipes has several optimized features that adapt it to large scale data analysis, namely:

- **Multiple schedulers**: GenPipes is optimized for HPC processing. It currently accommodates 4 schedulers.
- **Job dependencies**: GenPipes establishes dependencies among its different steps. This enables launching all the steps at the same time and minimizes queue waiting time and management.
- **Smart relaunch**: GenPipes sets and detects flags at each successful step in the pipeline. This allows the detection of successfully completed steps and easy relaunch of failed steps.
- **Parameter encapsulation**: Genpipes uses a superposed configuration system to parse all required parameters from configuration files. This simplifies the use of the framework and makes it more flexible to user adjustments. Tested configuration files that are tailored to different clusters and different species are included with GenPipes.
- **Diverse inputs**: GenPipes has been developed to launch using different starting inputs, making it more flexible.
- **Flexible workflows**: GenPipes implements a workflow in steps. Users can choose to run specific steps of interest, limiting waste of time and resources.

GenPipes is under continuous development to update established pipelines and to create new pipelines for emerging technologies. For instance, new genomics pipelines are being developed for ATAC-Seq, single cell RNA-Seq and HiChIP. GenPipes is also being redeveloped to use the Common Workflow Language (CWL) to provide a cloud compatible version more seamlessly and more *Scheduler* objects, like DRMAA, are being added to expand compatibility with more platforms. GenPipes has become a reliable bioinformatics solution that has been used in various genomics publications for DNA-Seq [60–67], RNA-Seq [68] and ChIP-Seq [69] analyses. GenPipes is currently available as source code, as well as a Docker image for easy installation and use. GenPipes has been optimized for HPC systems but can run on a laptop computer on small datasets.

## Availability and requirements

- Project name: GenPipes
- Project home page: http://www.c3g.ca/genpipes
- Operating system(s): Linux; Can be used on Windows and Mac OS using Docker
- Programming language: Python
- Other requirements: Workflow-dependant; detailed in documentation
- License: GNU GPLv3
- SciCrunch RRID: SCR_016376

## SUPPLEMENTARY DATA

No Supplementary Data

## ACKNOWLEDGEMENT

Data analyses were enabled by compute and storage resources provided by Compute Canada and Calcul Québec. Authors would also like to acknowledge Romain Gregoire and Tushar Dubey for their contribution to the code and Patricia Goerner-Potvin for her help in planning the report content.

---

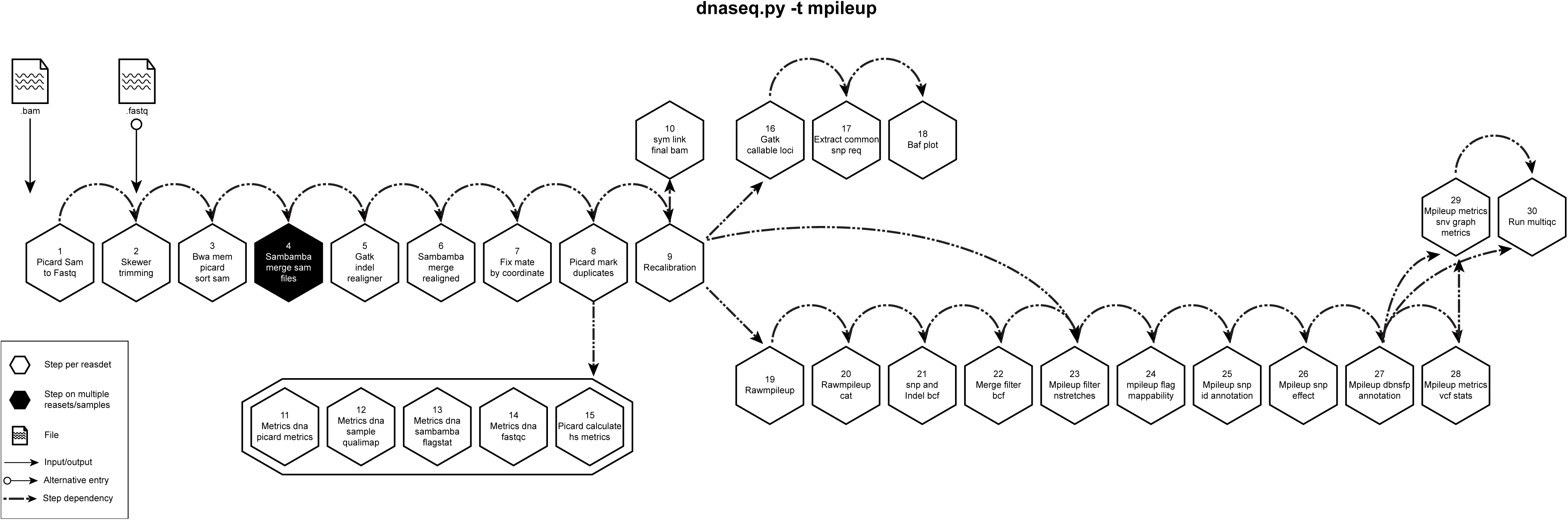

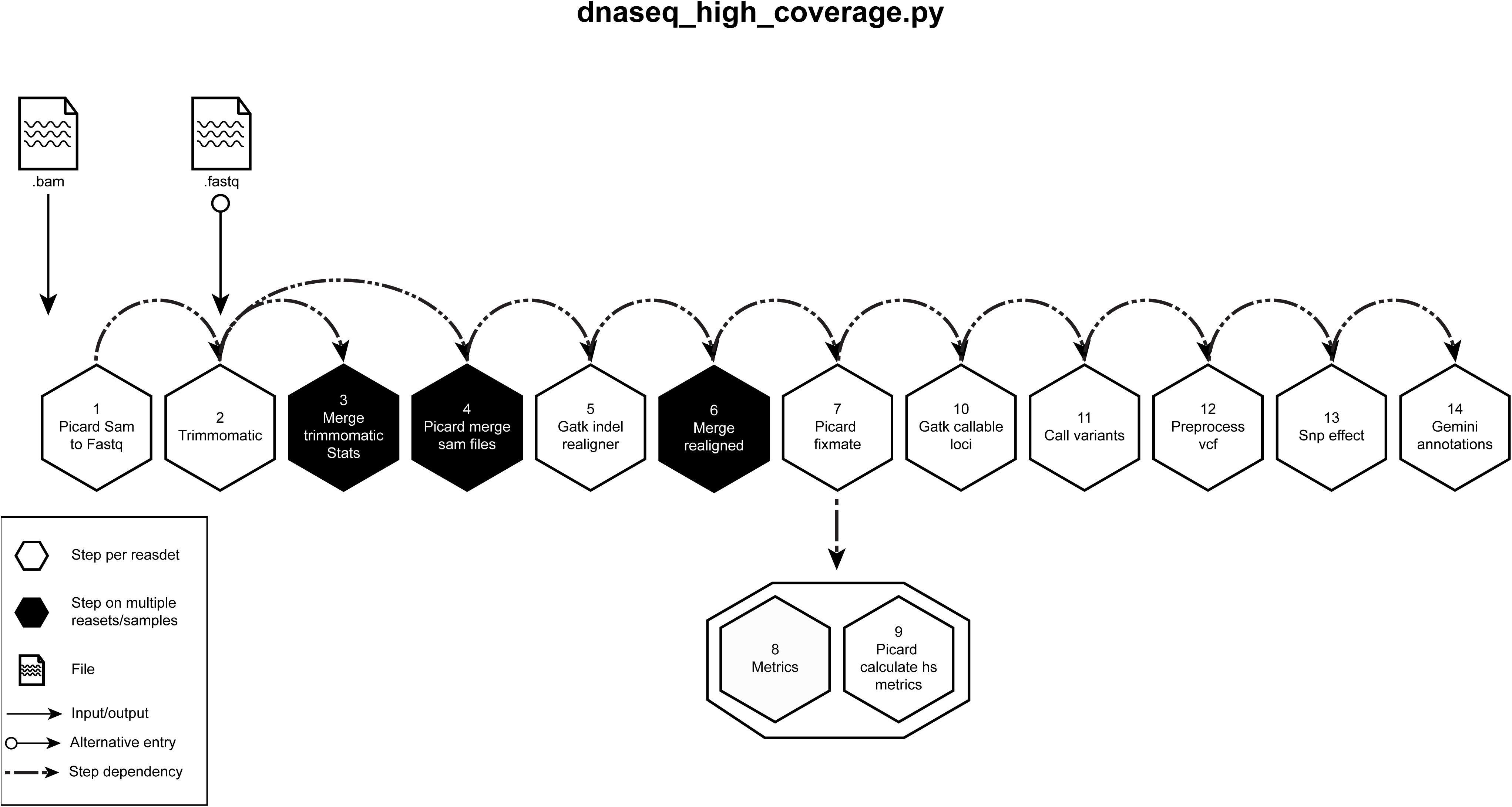

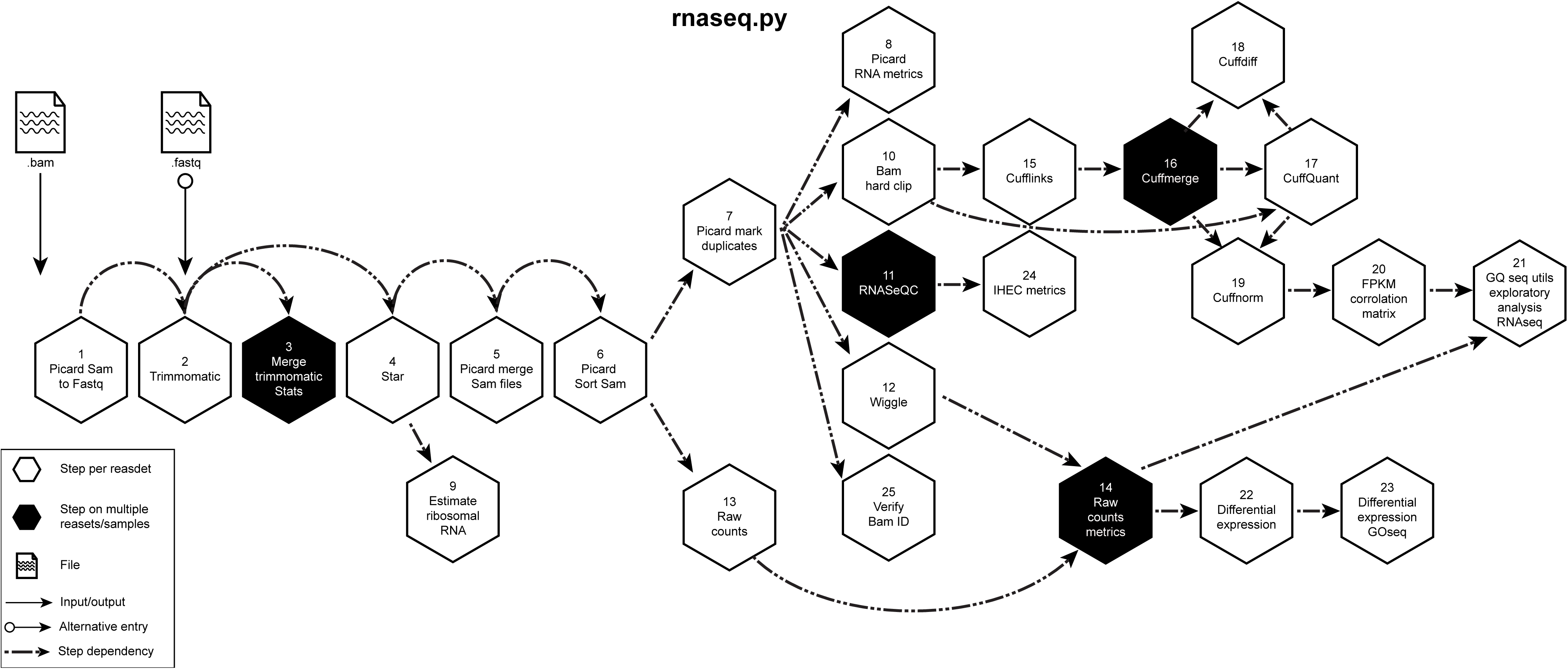

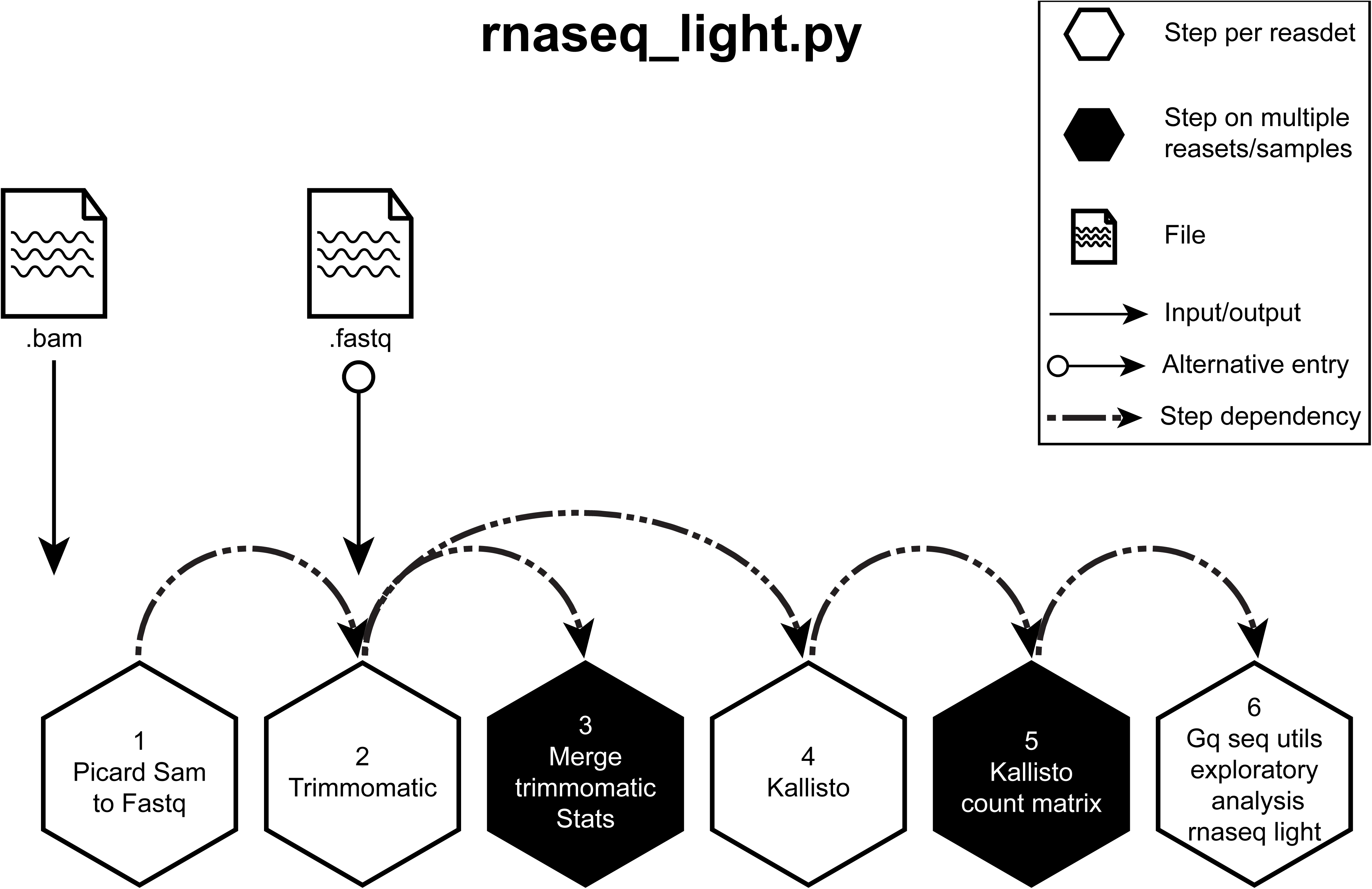

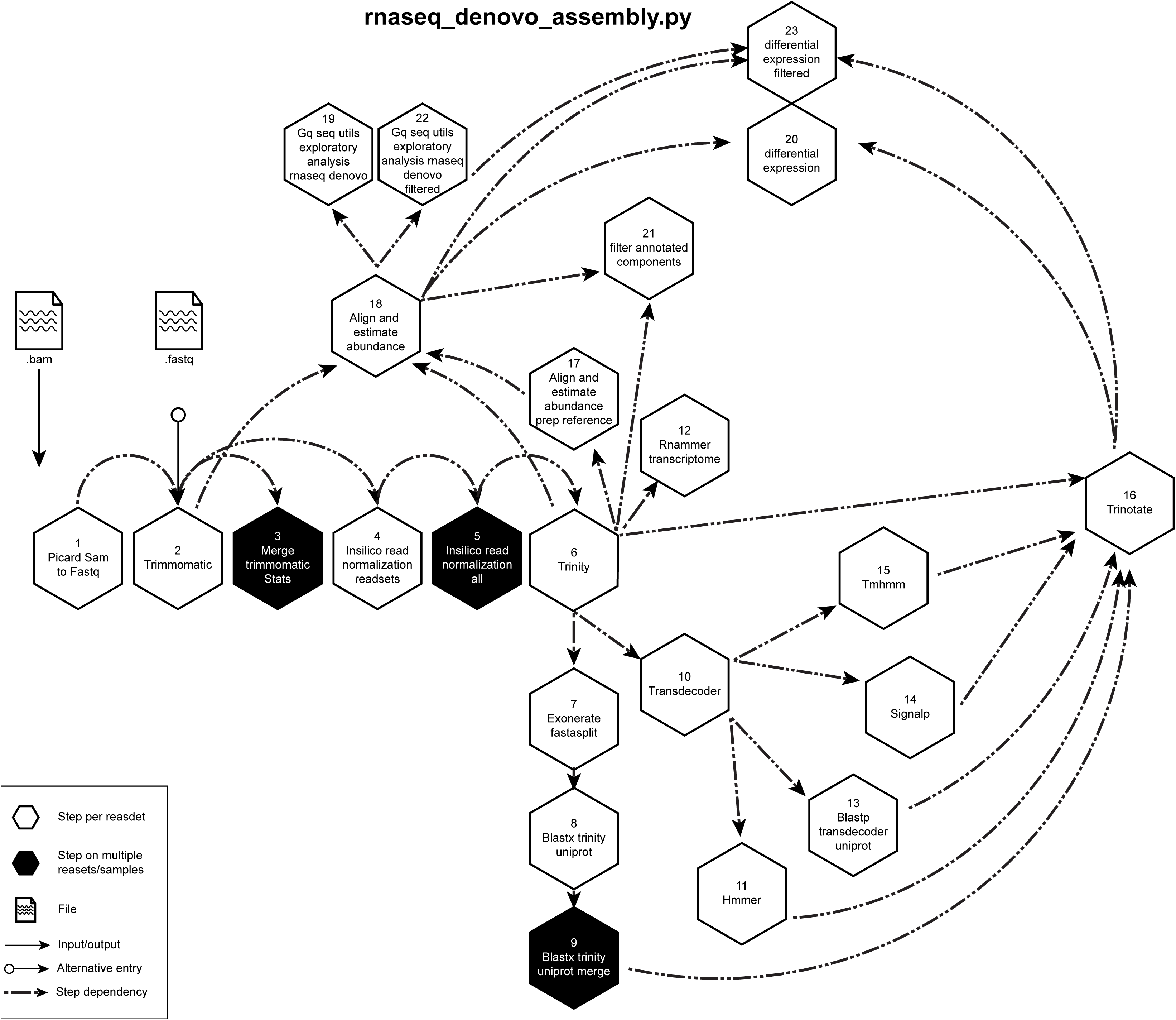

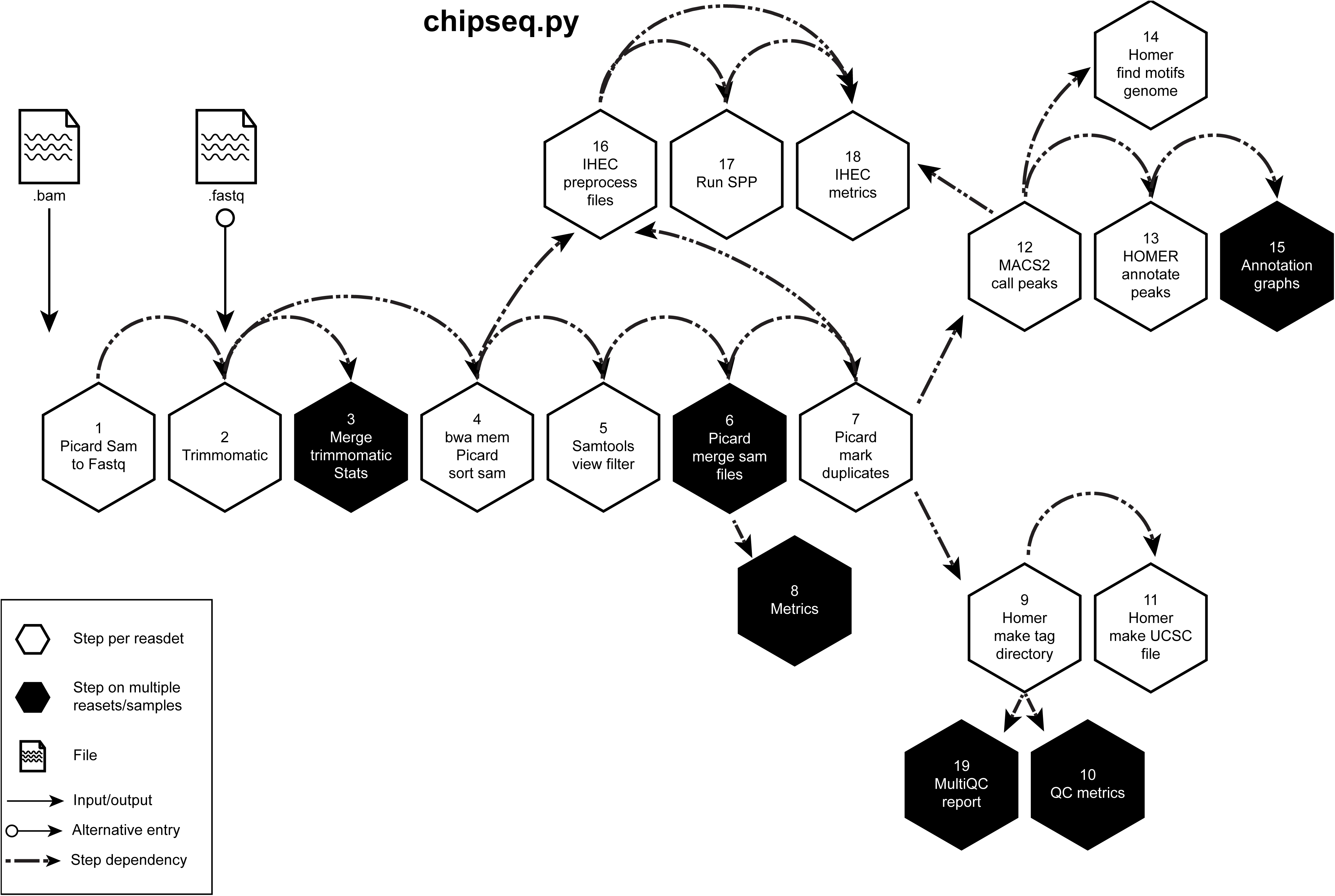

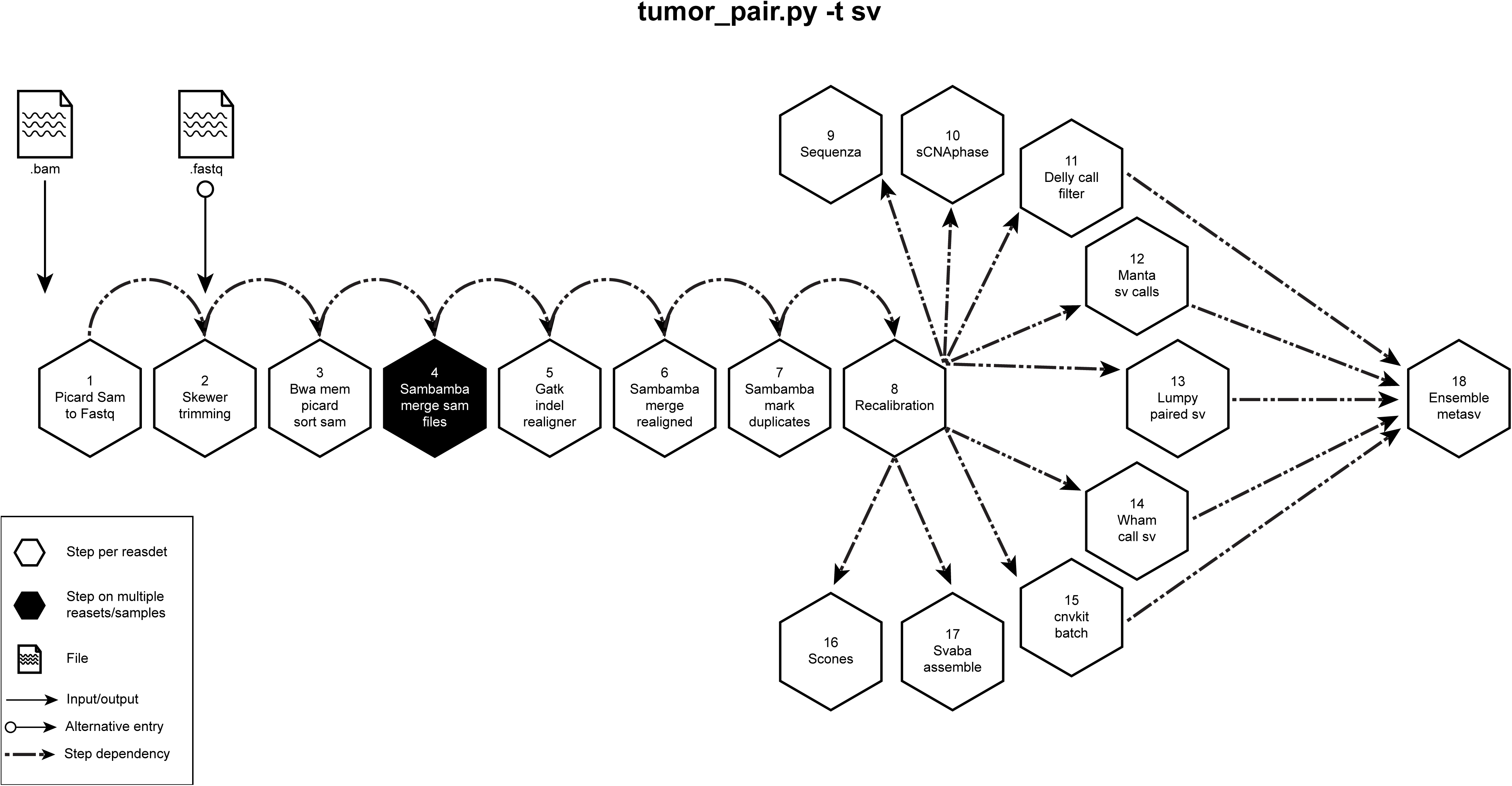

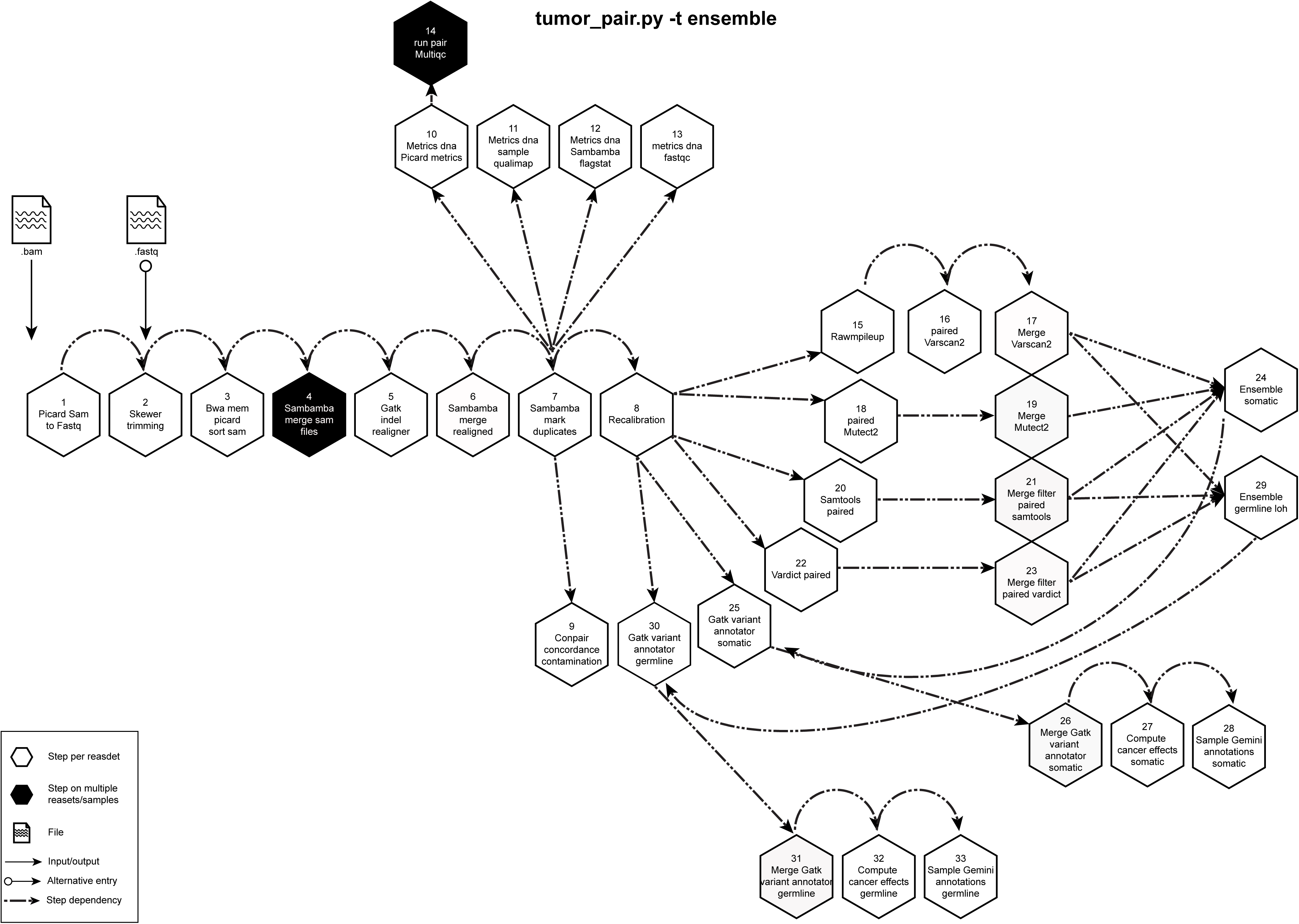

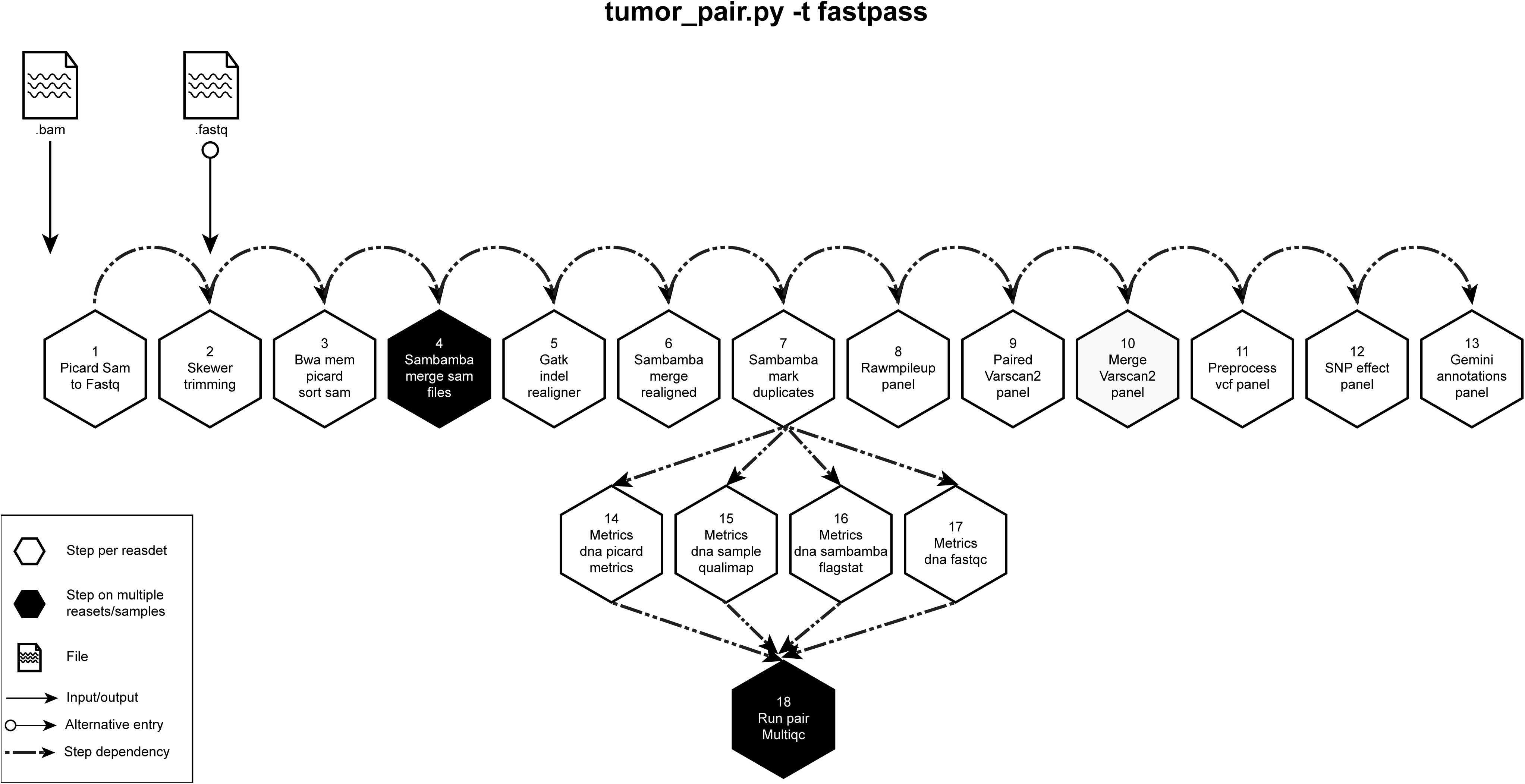

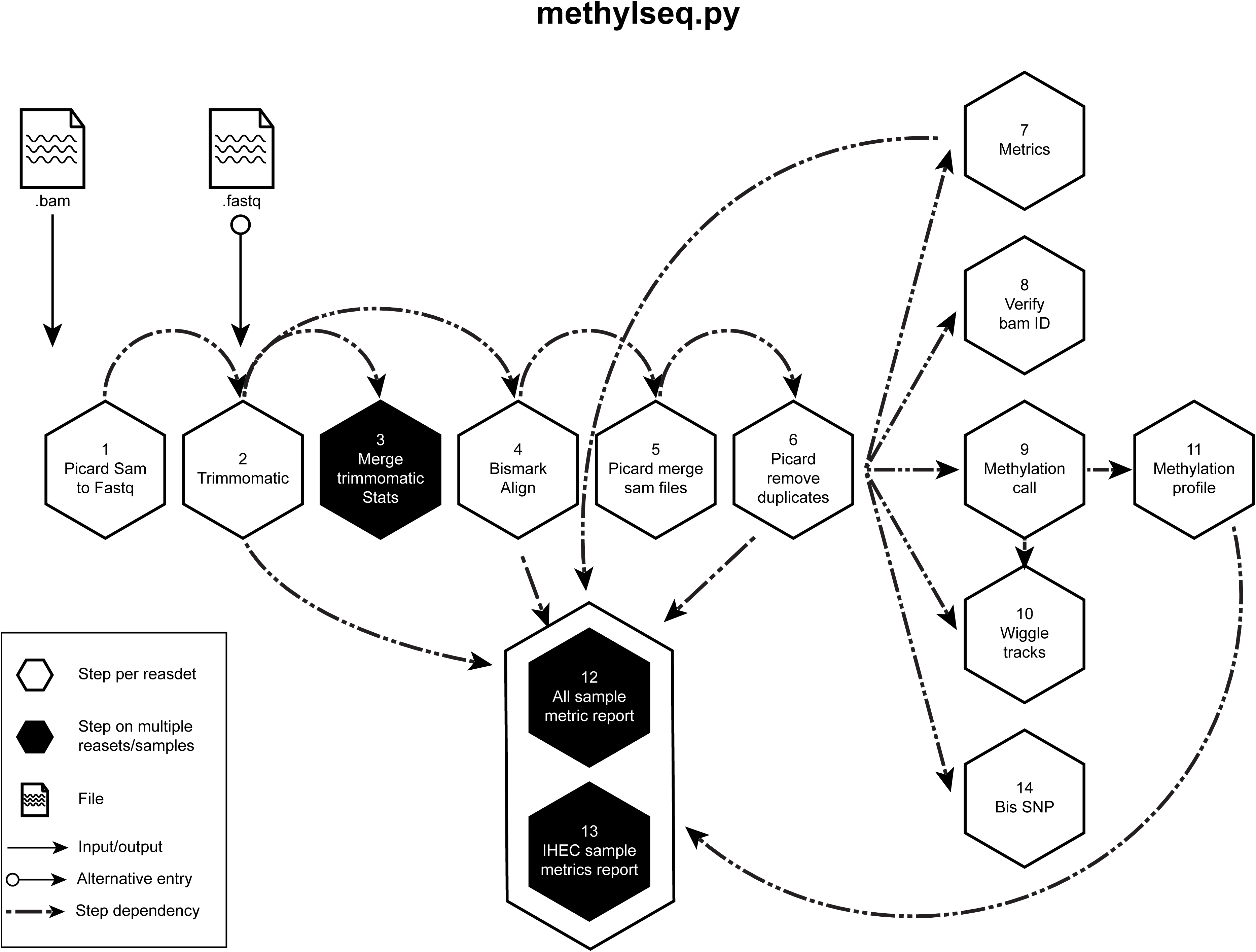

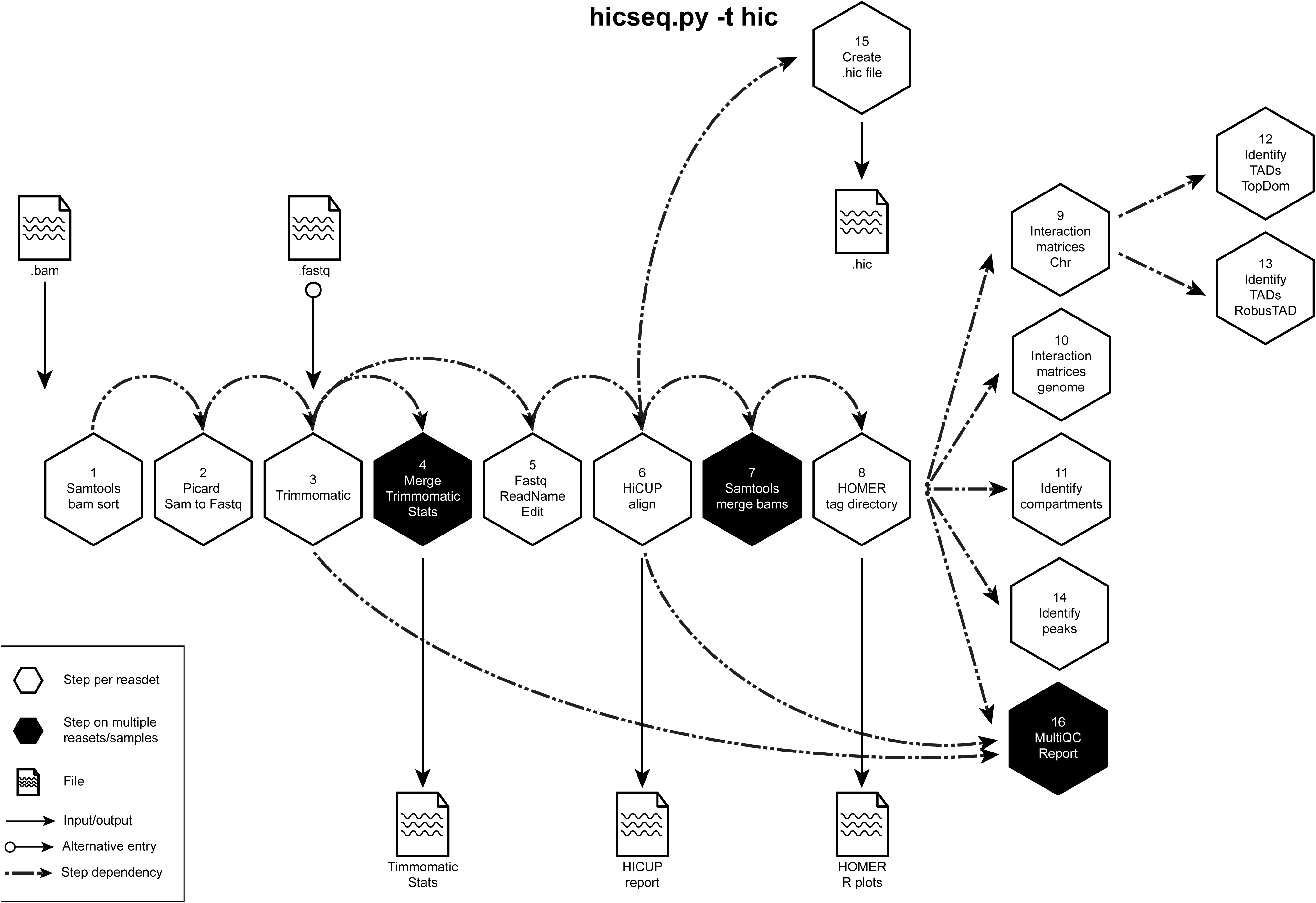

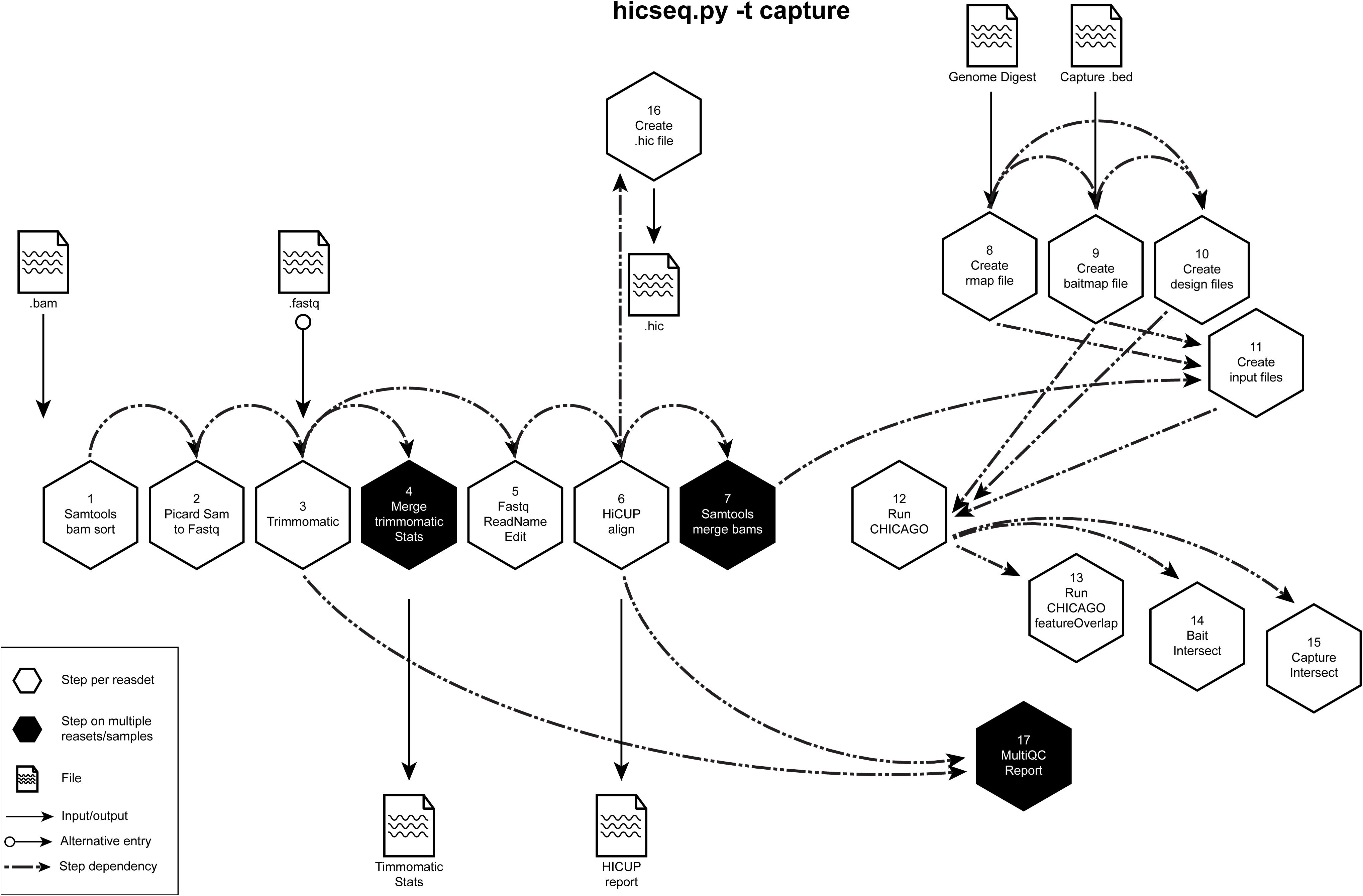

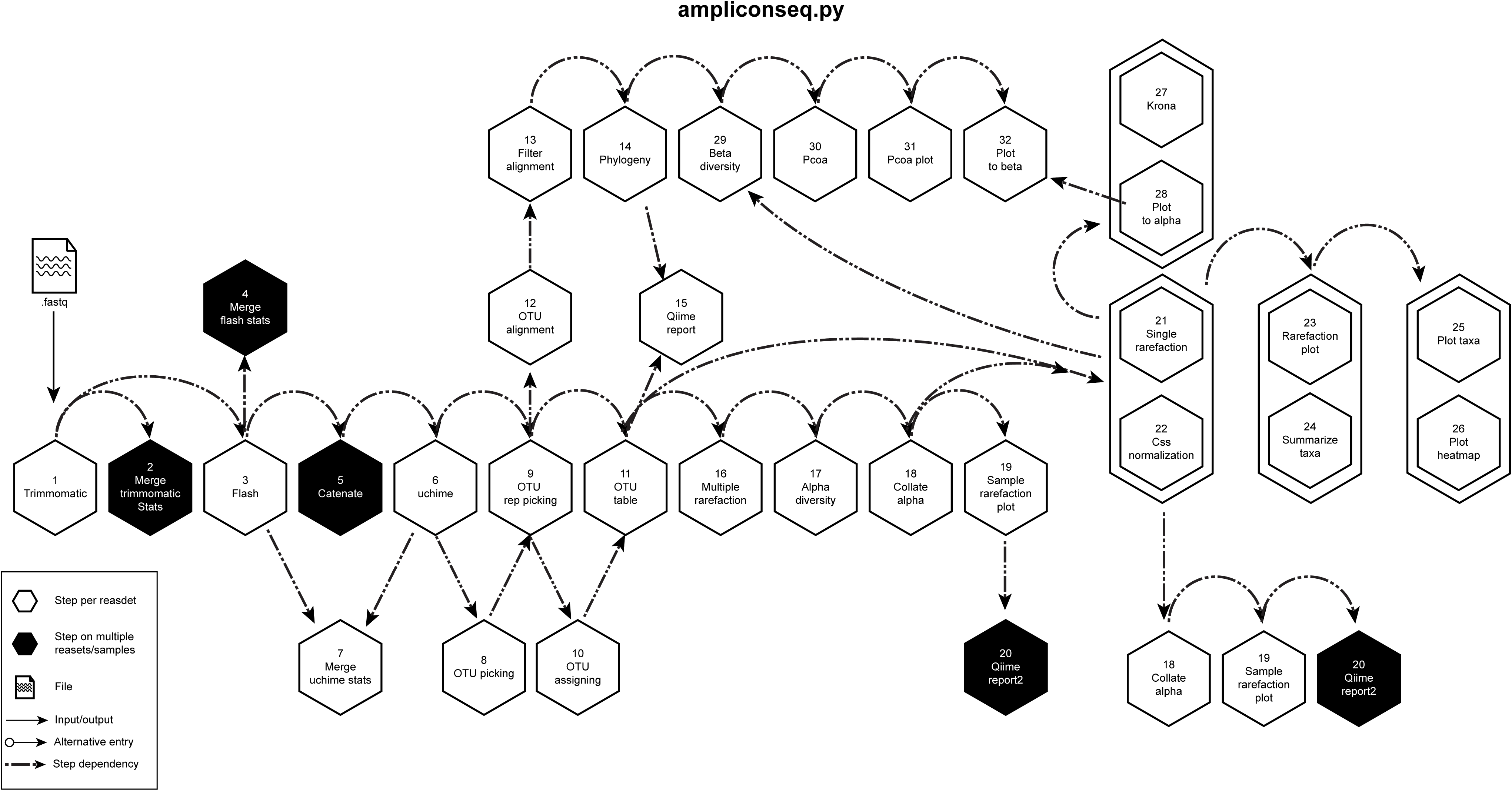

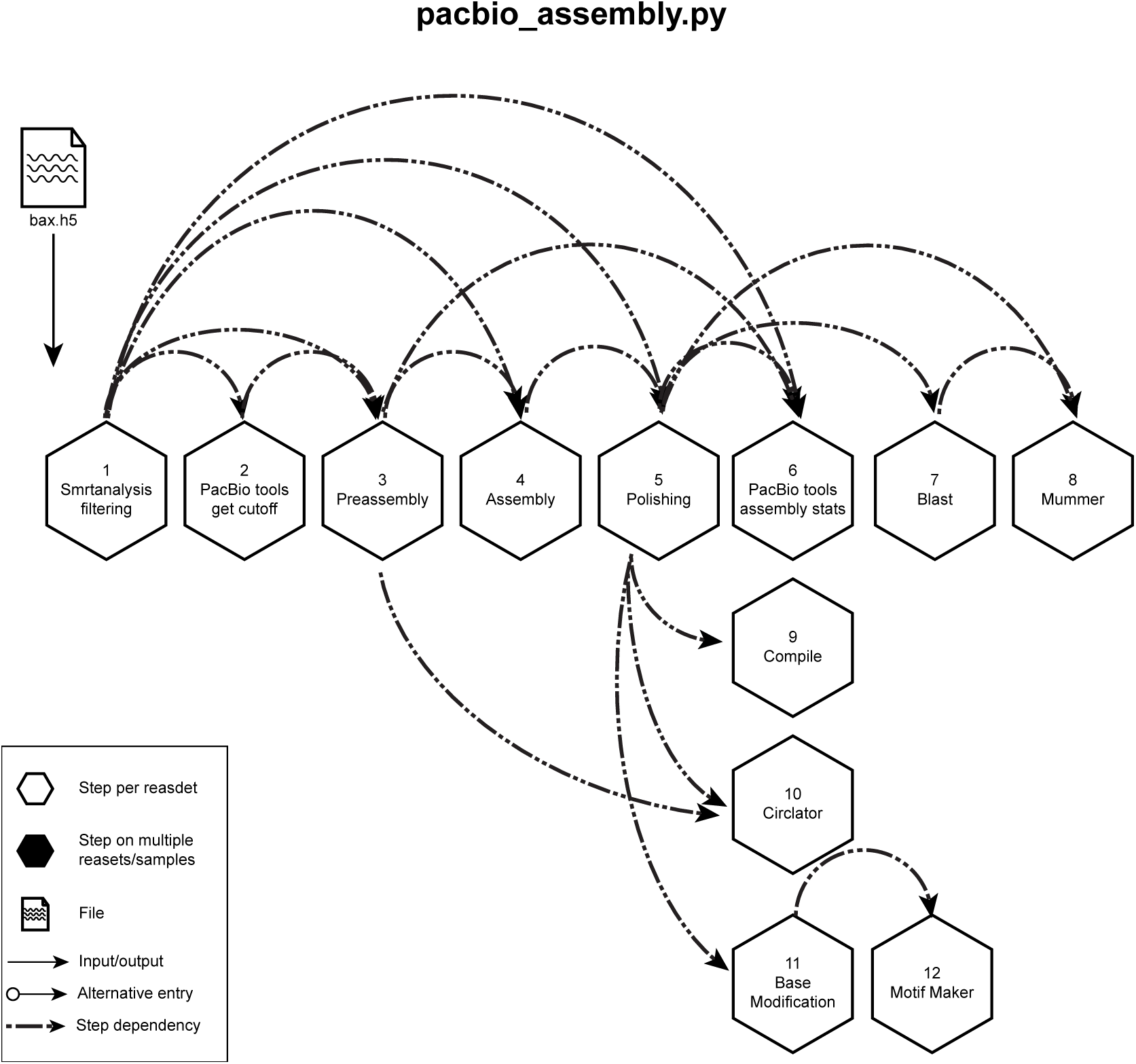

